# Perinatal Inflammation Influences but Does Not Arrest Rapid Immune Development in Preterm Babies

**DOI:** 10.1101/720888

**Authors:** S. Kamdar, R. Hutchinson, A. Laing, F. Stacey, K. Ansbro, M.R. Millar, K. Costeloe, W.G. Wade, P. Fleming, D. L. Gibbons

**Author notes:** Correspondence should be addressed to; 020 7848 9608 (phone). these authors contributed equally to the paper (P. Fleming, D. L. Gibbons).

## Abstract

Infection and infection-related complications are important causes of death and morbidity following preterm birth. Despite this, there is limited understanding of the development of the immune system in those born prematurely and how it is influenced by perinatal factors. To investigate this, we prospectively and longitudinally followed a cohort of babies born before 32 weeks of gestation. We demonstrated that preterm babies, including those born extremely prematurely, were capable of rapidly acquiring adult levels of immune functionality; that immune maturation appeared to occur independently of the developing microbiome, which was highly heterogeneous across different infants; and that the biggest drivers of change in the trajectory of perinatal immune development was exposure to infection *in utero* or post-natally. Conspicuously, a unifying factor among infants who developed infection despite their growing immune potentials was an inability to mount adequate T cell CXCL8 responses. Because this defect was present at birth, it may predict those babies likely to have poor clinical outcomes.

## 1 Introduction

Longitudinal data consistently show overall mortality following preterm birth to be falling, however among babies who die the proportion attributed to infection and infection-related complication such as necrotising enterocolitis is fixed or increasing ^1, 2^. It is estimated that up to 30% of preterm babies born before 32 weeks’ gestation develop at least one clinical or microbiologically confirmed episode of sepsis ^3^, but the course of individual babies through the perinatal period is unpredictable, some progressing smoothly while others succumb to multiple pathologies including late-onset sepsis and necrotising enterocolitis.

There is clinical need both to identify around the time of birth those preterm infants most likely to develop infective complications and to develop therapies to prevent and ameliorate infection in this vulnerable group, ideally through promoting the baby’s own immune capabilities. Understanding the development of the immune system in the healthy preterm infant, how it differs in those with complicated courses and how it is modified by peri and postnatal factors is fundamental to these aims. Furthermore, not only are these issues acutely relevant to the infant, but increasing evidence suggests that the nature of the infant immune response in early life, where rapid immune development has been observed ^4, 5^, may profoundly influence later susceptibility to immune mediated diseases ^6, 7^. Thus, infants born before 32 weeks’ gestational age [GA] show a greater susceptibility to asthma in later life and yet a decreased risk of developing food allergy ^8,9^.

Despite these drivers, understanding of how the preterm immune system develops is incomplete and the impacts on the developing immune system of exposure to infection *in utero*, and the postnatal clinical course have not been comprehensively evaluated.

It is increasingly acknowledged that the preterm immune system is not just an immature version of the adult but is both qualitatively and quantitatively distinct with overt functional potentials ^10^. To address how the immune system develops and to assess the impacts upon this of preterm birth, we prospectively followed a cohort of preterm babies born before 32 weeks’ gestation and compared their immune function over time. During the three months we studied this cohort, we demonstrate that preterm babies (even those born at the earliest gestation) rapidly acquire some adult levels of immune function; have immune functions that progress towards but do not reach adult levels; or have immune functions, conspicuously CXCL8-producing T cells, that remain significantly distinct from adults. Babies born at an earlier GA showed immune profiles that changed more dramatically when compared to those born at later gestation, thus reducing any negative impact of extreme prematurity. Immune development occurred independently of specific changes within the developing microbiome, suggesting other factors have a greater bearing on immune development in the first few months of life. A defining feature of those infants that did develop infection was a lower level of CXCL8-producing T cells at birth thus identifying a potential biomarker for infants at greater risk of sepsis.

## 2 Methods

### 2.1 Ethical Approval and Funding

Funding was granted by Barts Charity (Ref: 764/2306) for the undertaking of a study named “Investigating Microbial Colonisation and Immune Conditioning in Preterm Babies”. Ethical and regulatory approvals were granted by the London (Chelsea) Research Ethics Committee (Ref: 15/LO/1924) and the local Research and Development department at Homerton University Hospital NHS Foundation Trust.

### 2.2 Participant Enrolment

Subjects were recruited from the neonatal intensive care unit at Homerton University Hospital between January 2016 until December 2017. Babies born between 23 weeks and 4 days and 31 weeks and 6 days of gestation admitted to the NICU before 72 hours’ postnatal age, including those born at other hospitals, were included in this study. Infants with a probable lethal congenital abnormality, thought to have no realistic chance of survival, or with known exposure to either HIV or hepatitis B infection were excluded. Written informed consent was obtained from parents before 72 hours of age.

### 2.3 Sample collection and storage

Blood samples (0.5 ml whole blood) were collected on a weekly basis, with additional samples taken at times of suspected infection, up to cessation of the subjects’ participation. These were stored at room temperature (on average for 12 hrs and up to a maximum of 30 hrs) prior to Ficoll separation of PBMCs, and were frozen in Cryostore (Sigma) and stored in liquid nitrogen. Faecal samples were collected on a daily basis (when available) from birth up to cessation of the subjects’ participation in the study (*i.e.* discharge or transfer). Samples were immediately stored at 4**°**C for <72hrs, before transferral to long-term storage at −80°C.

### 2.4 Clinical data

Detailed demographic, antenatal and daily postnatal data were collected prospectively. Once each subject completed their hospital stay, their clinical course was reviewed by the clinical team and the subjects classified *a priori* into one three clinical cohorts:

- Stable infants
  -infants receiving a single course of antibiotics only in the immediate postnatal period, with no episodes of suspected late onset sepsis or antibiotic exposure thereafter
- Unstable infants
  -infants who did not meet the stable criteria, having received additional courses of antibiotics for suspected late-onset sepsis
  -Unstable infants were subsequently divided into those with, or without, a history of clinical or histologically confirmed chorioamnionitis.
- The collaborating immunology team was not aware of the clinical status of babies when blood samples were collected and/or analysed. Once the analyses were completed, the subjects’ clinical courses were provided.

### 2.5 Cell stimulation

Peripheral blood mononuclear cells were stimulated with Phorbol Myristate Acetate (PMA) and ionomycin as described below. Cells were stimulated in RPMI 1640 medium (Invitrogen) containing 10% (vol/vol) FCS (StemCell Technologies), 2 mM L-glutamine (Sigma), 100 U penicillin and 100 μg/ml streptomycin (Invitrogen) (complete medium, CM) containing 10 ng/ml phorbol 12-myristate 13-acetate (Sigma) and 1 μg/ml ionomycin (Sigma) for 4 h at 37°C, 5% CO2 in the presence of 20 μg/ml brefeldin A (Sigma). For spontaneous cytokine production, cells were incubated in CM in the presence of brefeldin A only.

### 2.6 Flow cytometry

All samples from individual babies were analysed simultaneously to avoid introducing any unnecessary experimental variation. Single-cell suspensions were prepared in FACS buffer (PBS plus 2.5% (vol/vol) FCS and 2 mM EDTA) and then split into 9 aliquots for staining with 9 different 11-colour flow cytometry panels. To one of the aliquots, 50 μl of Countbright Absolute counting beads (Catalogue no.C36950, Life Technologies) was added. Initially, cells were incubated on ice for 15 minutes with Zombie NIR Fixable viability kit (Biolegend) to allow live/dead discrimination. For panels containing Vδ1, the surface staining antibody was added along with the live/dead discrimination stain. Cells were then surface stained on ice for 30 min with antibodies listed in **Supplementary Table 1**. Following surface staining, cells were fixed (Cell Fix, BD) and then permeabilised using Perm buffer (BioLegend). For panels containing transcription factor FoxP3, cells were fixed and permeablilised using fixation and permeabilising reagents from Invitrogen’s Intracellular Fixation & Permeabilisation Buffer Set (Catalogue no. 88-8824-00) according to the manufacturer’s instructions. Following permeabilisation, cells were then intracellulary stained for cytokines and transcription factor FoxP3 with antibodies listed in **Supplementary Table 1**. All samples were run on a 4 laser BD LSRFortessa X20 flow cytometer. To minimize impact of temporal variation in instrument sensitivity, cytometer performance was monitored using CS+T beads (BD) and experiments were run using applications settings. Data were analysed using FlowJo software.

### 2.7 Laboratory methodology for faecal DNA extraction

Faecal samples were selected for analysis on a pragmatic basis to provide regularity of assessment throughout the subjects’ admissions, with increased resolution at times of clinical instability; and subject to availability – samples were taken approximately every three days (where possible), with additional samples taken before/during/after courses of antibiotics, feeding type changes, ventilation changes, and blood transfusions, to capture possible associated microbiome changes.

Faecal bacterial DNA was extracted using the DNeasy®PowerSoil®Kit (Qiagen, Netherlands), following treatment with propidium monoazide, to prevent downstream amplification of contaminant or non-viable free bacterial DNA. The V4 hypervariable region of the 16S rRNA gene was amplified using PCR with a dual-indexing strategy, and sequencing of the subsequent amplicons was performed using Illumina MiSeq technology (2x 250 bp flow-cell for paired-end sequencing with 5% PhiX DNA).

### 2.8 Faecal microbiome analysis

Sequencing data were processed using the DADA2 pipeline and all analyses and graphics were produced by means of R packages (DADA2, decontam, tidyr, dplyr, ggplot2 and mblm) ^11 12 13 14-17^. Microbiome analyses were performed on 10 stable (100% of eligible subjects), 10 unstable (63%) and 11 unstable chorioamnionitis (84%) subjects.

Biometrics of microbiome structure were used to describe the development of bacterial communities within the cohort: for the description of constituent taxa, analysis was performed at family-level, as this represented the lowest taxonomic level with the highest level of read identification (99.9% of amplicons were identified at family level; only 74% were identified at genus level); only those taxa with a mean relative abundance of >1% across all samples were included in the analyses. Diversity was estimated using the inverse Simpson statistic, and was derived from all taxa (regardless of relative abundance). The longitudinal progression of the relative abundances of individual family-level taxa were described.

### 2.9 Statistical Analysis

#### Longitudinally changing datasets

To identify immune parameters that changed significantly over time, longitudinal models of the cell-subset frequencies and total numbers at the group level were constructed. Group level slopes were estimated using mixed linear models of the cellular frequencies/numbers versus post-natal age, with a random intercept and slope at an individual level (LME4 R package) ^18^. The regression lines were weighted by the number of samples/per baby which varied between subjects. To determine whether a given immune parameter changed significantly between one timepoint and another, a nonparametric Wilcoxon matched-pairs signed rank test was used.

To assess the longitudinal development of microbial communities over time, regression coefficients (Theil-Sen non-parametric regression coefficient) for diversity and *Enterobacteriaceae* relative abundance against time were calculated as summary measures to describe the longitudinal progression of individual subjects. These summary measures were then used to derive group level summary statistics (*i.e.* median), and to allow statistical comparison between the groups [by weighted Mann-Whitney U-test]. Summary measures were weighted to account for the number of originating data points (corresponding to numbers of samples) from which the individual measures were derived. Only the family *Enterobacteriaceae* was examined in this manner, as it was the only taxon widely distributed across the data set in significant relative abundances (>40% mean abundance in >90% of all subjects) to allow comparison between the three clinical groups. Longitudinal progression of individual taxa in the three clinical cohorts is represented by a plot of relative abundance of individual taxa at each time point. In recognition of the bimodal distribution of some taxa, we highlighted the subjects who were essentially not colonised with specific taxa at any time point (maximum relative abundance <1%).

To understand the relationships that exist between immune parameters and microbiome parameters, intra-individual Spearman correlation was performed. To account for variation in the sampling schedule between blood and stool we binned the data into 5-day time windows to match blood and contemporaneous stool samples. In the event of more than one sample per individual occurring in any given bin, mean values were calculated for every parameter measured. Spearman correlation analyses were performed across all parameters barring those directly linked to one another owing to the nested nature of flow cytometry analysis. A mean value was derived for the correlation coefficient between all parameters across all individuals. For a relationship to be considered significant it would have to consistently yield a p-value of <0.2 in every individual tested, the aggregate of this across all individuals equates to a p value threshold of p=0.008. Stool passage was unpredictable among babies recruited to the study so not all infants had sufficient samples to perform a 100% match between immunological and microbiome parameters.

## Supporting information

All Supplementary data

## 2.10 Acknowledgements and Author contributions

We thank S. Evans for help with statistical analyses; I. Jogee for illustrations; A. Das and F. Kyle for some sample processing; and A Hayday (KCL) for critical review of the manuscript. PF, RH, FS, KA, KC and MM were all supported by a strategic research grant from Barts Charity; SK and DG by Cancer Research UK (grant A20730); AL by the Wellcome Trust (grant 100156/Z/12/Z to A Hayday), and WGW by Queen Mary University of London.

PF, RH and FS did all the patient and sample recruitment; SK did blood processing, all the flow cytometry experiments and data analysis; RH and KA did DNA extraction from stool, Next Generation Sequencing and RH analysed the microbiome data; AL helped design and optimise flow cytometry panels and analysed large data sets; WGW, MM and KC provided critical review of the data; PF set up the study, evaluated the clinical data and co-wrote the manuscript; DG designed the immunology study, evaluated the results and co-wrote the manuscript.

There are no competing interests to declare.

## 3 Results

### 3.1 Patient Demographic and Clinical Characteristics

Thirty-nine babies with a median gestation of 28.7 weeks (IQR 25.7-30.4) and a mean (SD) birthweight of 1060g (383) were enrolled between January 2016 until December 2017. Participant demographic and clinical data are provided in **Table 1**. Ten babies were classified as stable (receiving only the standard single course of antibiotics in the immediate postnatal period) and 29 were unstable, having received additional courses of antibiotics for suspected late-onset sepsis. Of the 29 unstable babies, 13 were born to mothers with histological or clinically confirmed chorioamnionitis. Of note, babies in the stable group were generally born at later gestation, had higher birth weights and all were delivered by caesarean section.

**Table 1:**
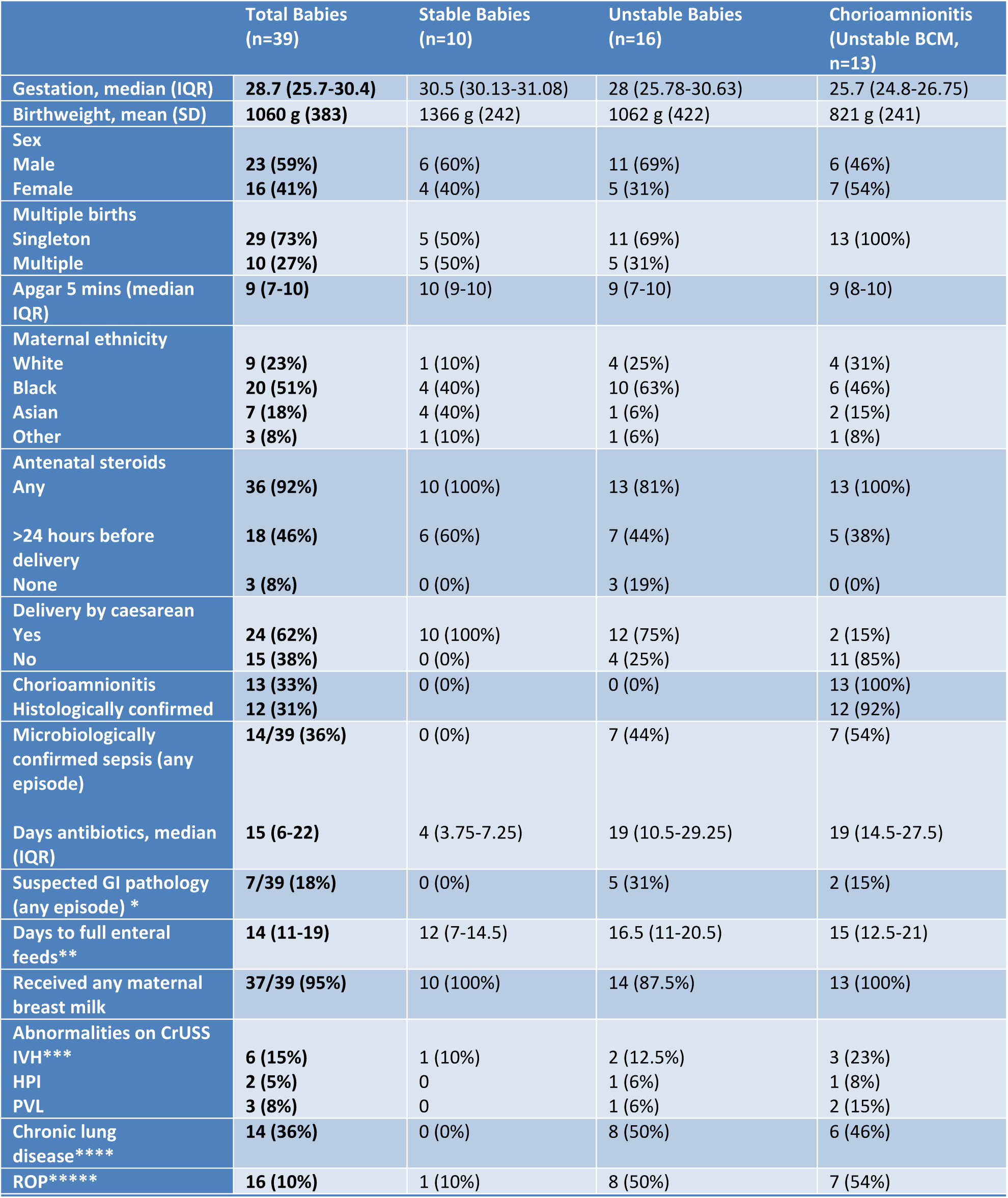
Data are presented as means (standard deviation); medians (interquartile ranges); n=number and percentages. *Suspected GI pathology is defined as ‘*any abdominal concerns necessitating nil by mouth for more than 5 days*’ **Full enteral feeds defined *as ‘the first day on which 100*% *of fluid volume was administered enterally*’; CRUSS=cranial ultrasound; ***IVH= intraventricular haemorrhage defined *as*’ *bleeding contained within and not extending beyond the ventricular system*’; HPI= haemorrhagic parenchymal infarction; PVL=periventricular leukomalacia; ****CLD= chronic lung disease defined as’ *the need for supplemental oxygen at 36 weeks postmenstrual age’*; *****ROP=retinopathy of prematurity defined as *‘any stage ROP recorded on formal eye examination*’

### 3.2 Postnatal Immune Development in All Babies

186 different peripheral blood immune populations were analysed, with and without *in vitro* stimulation, by use of 11-colour flow cytometry panels. Samples were taken longitudinally (mean of 8 samples/baby) from the infant cohort until discharge from the NICU or until 36 weeks’ postmenstrual age. The sampling regime, analysis pipeline and clinical groupings are depicted in **Supplementary Fig 1**. Initially we identified immune parameters that were most distinct between adults and preterm infants at birth (or within the first week of birth) (**Supplementary Fig 2)**. Unsurprisingly, CXCL8 (a cytokine known to be pivotal in this demographic ^10^) was one of the most divergent parameters, significantly elevated in infants compared to adults.

#### Changes in immune parameters over time

We then studied any alterations in these immune parameters over time. To address which parameters changed most significantly with postnatal age, we compared the regression coefficients of each parameter in each baby and plotted the mean value in relation to how different they were from adult at birth (**Fig 1A**). Over half of all parameters studied showed significant changes in this time period. Many parameters that were reduced at birth (anything below the half way line on y axis) increased (and hence appeared in lower right hand quadrant), and conversely those that were elevated at birth (anything above the half way line on y axis), decreased towards adult levels -located in the upper left hand quadrant. Conspicuously, although CXCL8-producing NK and γδ cells that were elevated at birth subsequently decreased with time, this was not true for CXCL8-producing CD4 and CD8 T cells of various phenotypes. These populations started elevated compared to adults and remained so, highlighting their potential functional importance in the neonatal immune response.

**Figure 1:**
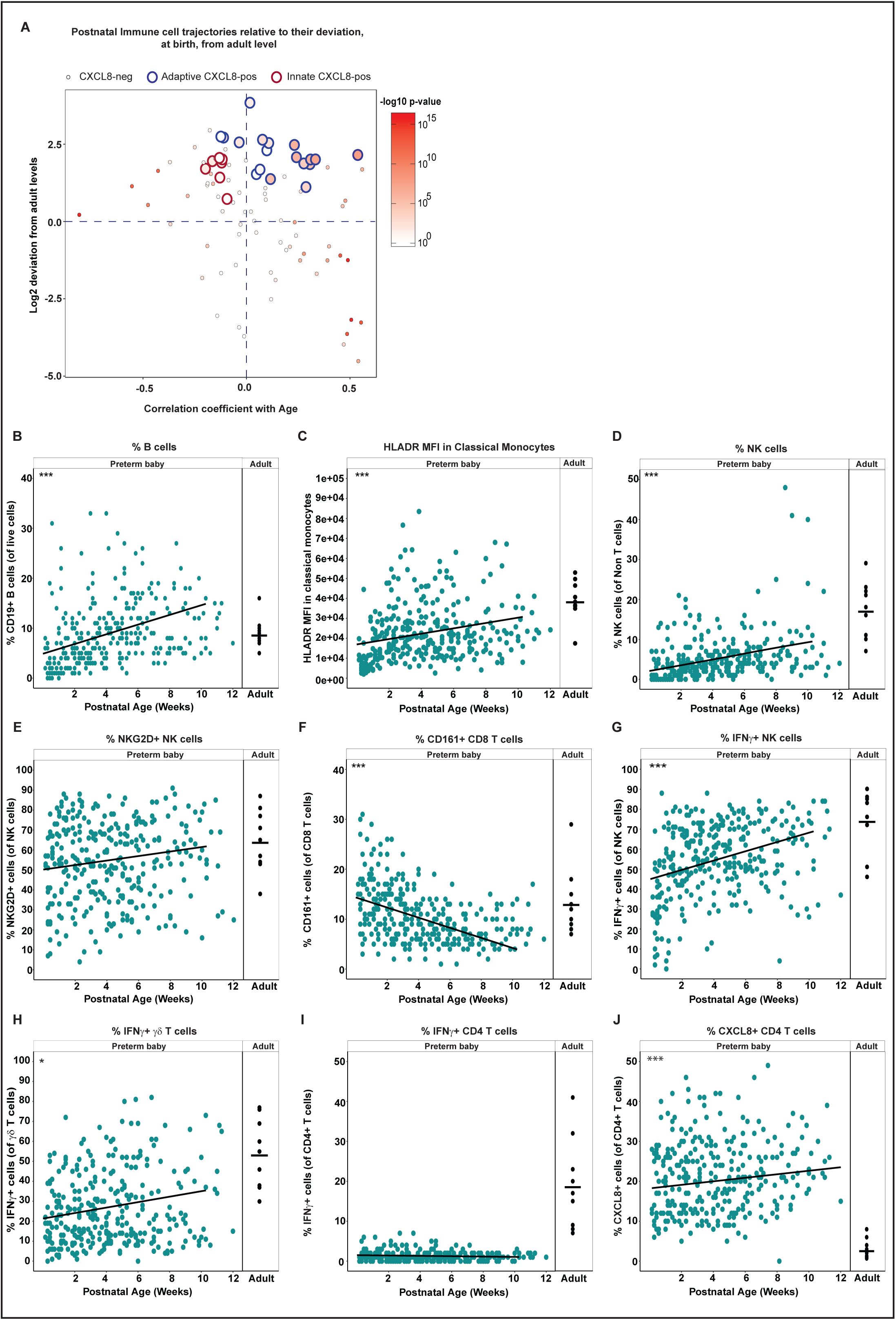
Phenotypic and functional maturation of distinct immune parameters. Longitudinal PBMC samples from 39 preterm babies were phenotyped for 186 different immune populations by flow cytometry following surface and intracellular staining. For cytokine detection, samples were activated in vitro with PI (4 h, in the presence of BFA) prior to staining. (A) Postnatal Immune cell trajectories relative to their deviation, at birth, from adult levels. Position along y axis indicates deviation from adult (the Log2 fold difference) at birth, position along x axis indicates whether the population has increased or decreased significantly in the infant over time. Thus, immune populations in the top left quadrant start higher than adult levels and then decrease over time towards the adult level. Populations in the bottom right start below adult levels and increase over time towards that of the adult. Colour indicates p value for correlation with age with large circles indicating CXCL8 producing cell types. Amongst the CXCL8-producing populations, the red outline indicates NK and γδ T cells whereas the blue outline CD4 or CD8 αβ T cells. (B-J) Changes in individual immune parameters over time depicted by scatter plots showing frequencies/MFI (as indicated) in preterm babies (left panel;cyan circles) as a function of postnatal age compared to adults (right panel; black circles) (B) CD19+ B cells, (C) HLADR MFI in classical monocytes, (D) NK cells, (E) NKG2D+ NK cells, (F) CD161+ CD8 T cells, (G) IFN-γ+ NK cells, (H) IFN-γ+ γδ T cells, (I) IFN-γ+ CD4 T cells and (J) CXCL8+ CD4 T cells. In the figure, cyan circles represent a pool of longitudinal samples from 39 preterm babies where each circle represents an individual sample and on average there are 8 longitudinal samples/baby. Black circles are a pool of samples from 9 adults. *** p< 0.001, **p < 0.01 and *p< 0.05 as determined by linear mixed effect modelling using the lmer package in R.

Parameters that increased over time included B cells that increased in both percentage (**Fig 1B**) and absolute number (**Supplementary Fig 3A**); HLADR expression on monocytes (**Fig 1C** and **Supplementary Fig 3B**); NK cells (**Fig 1D** [%] and **Supplementary Fig 3C** [absolute number]); and NKG2D expression on NK cells (**Fig 1E** [%] and **Supplementary Fig 3D** [absolute number]). Consistent with the observation that the CD4:CD8 ratio is much greater in (term) neonates than adults ^19^, the percent representation of CD8^+^ T cells also increased over time coincident with a decrease in CD4^+^ T cells (**Supplementary Fig 3E/F**). CD161 expression on CD8 T cells decreased in the 3-month study period (**Fig 1F**).

With respect to changes in functionality that occurred postnatally (determined by assessing cytokine production following *in vitro* stimulation with PMA and Ionomycin [P/I]), we observed significantly increased percentages of IFN-γ-producing NK cells [**Fig 1G**] and γδ T cells (**Fig 1H** and **Supplementary Fig 4A**). However, levels of IFN-γ-producing CD4 T cells in these babies remained extremely low even at 3 months of age and there was no reproducible increase in the capacity of CD4 T cells to produce IFN-γ over time (**Fig 1I** and **Supplementary Fig 4B**). CD4 T cells continued to produce CXCL8, however, and levels did not diminish towards that seen in adult in the time frame studied (**Fig 1J**).

#### Effect of extreme prematurity

To identify whether the consensus pattern of immune development that we observed in our infant cohort also occurred in the event of extreme prematurity, we performed PCA analyses using all the flow cytometry parameters to generate immune profiles for each baby at each time point. By comparing the immune profile generated in the first week of life compared to that generated at around 6 weeks of age, we observed that the babies’ immune profiles progressed in a similar direction as they aged (**Fig 2A**).

**Figure 2:**
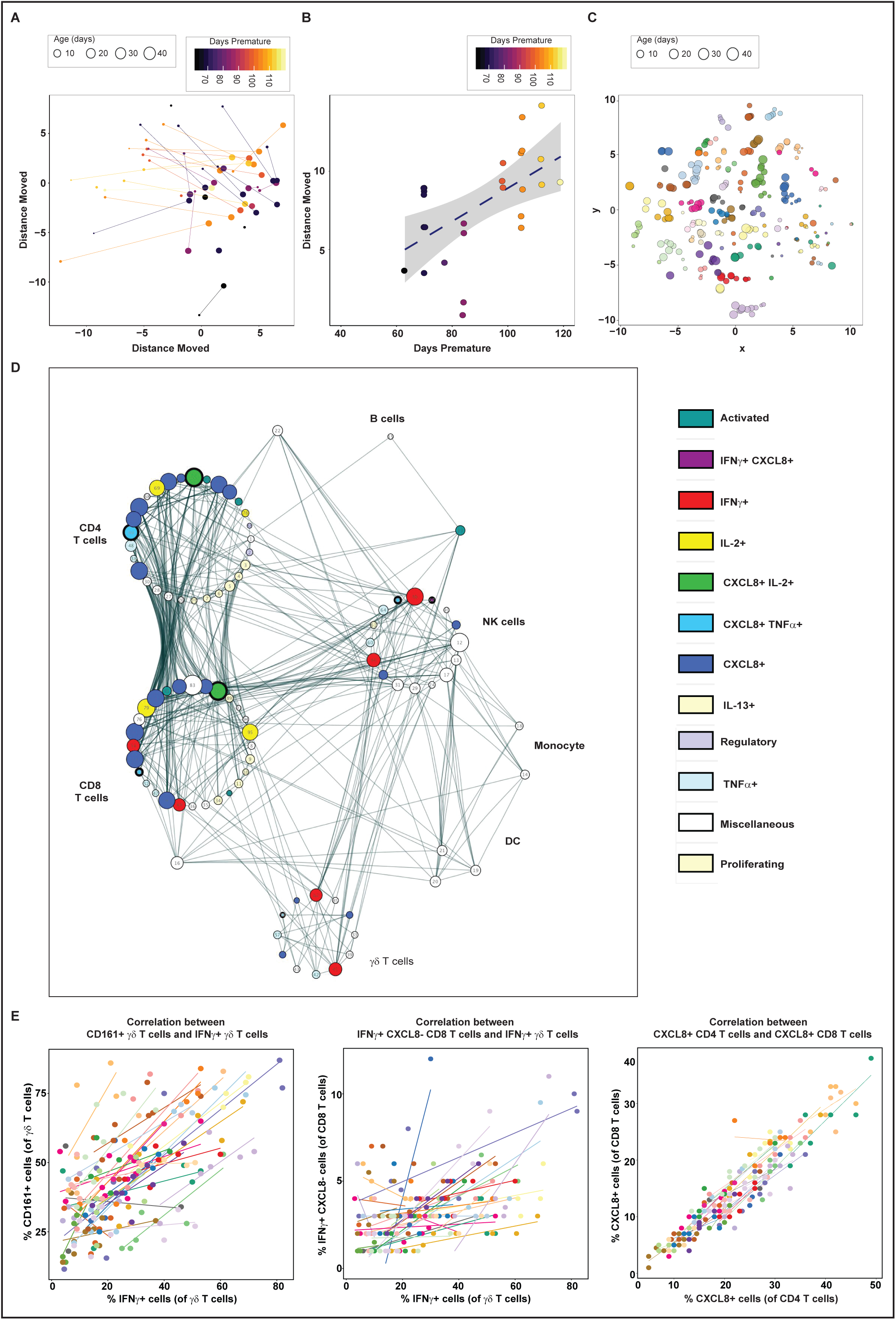
Development of whole immune profiles and correlations. Longitudinal PBMC samples from 39 preterm babies were phenotyped for 186 different immune populations by flow cytometry following surface and intracellular staining. For cytokine detection, samples were activated in vitro with PI (4 h, in the presence of BFA) prior to staining. (A and B) Analyses of the immune parameters from 39 preterm babies shows that the immune profile of extremely preterm babies (lighter colour) travels a longer distance but on the same trajectory compared to their less preterm counterparts (darker colour) to reach the common immune profile as depicted by: (A) PCA of the immune parameters showing the immune profile of the sample taken in the first week (small circle, mean=3.8days) to the immune profile of the sample taken 5 weeks later (large circle, mean=37days) for each baby. The colour gradient represents how many days preterm the baby was at birth (counting down from 40 weeks) as described in the figure. Each circle represents a sample from an individual baby and the size of the circle represents postnatal age of the baby at the time the sample was taken. Individual babies are linked by lines. (B) Scatter plot quantifying the Euclidian distance moved in PC1 and PC2 for each baby. The colour gradient represents how many days preterm the baby was at birth. Each symbol represents an individual baby. (C) tSNE analysis of the immune parameters shows that babies have their own individual profile that distinguishes them from each other. Each circle represents a sample from an individual baby (the size of the circle depicts the postnatal age of the baby when the sample was taken) and each colour represents longitudinal samples from the same baby. (D) Network represents statistically significant (p<0.008) spearman correlation coefficients (R=>0.3 or R=<-0.3) between immune parameters. Each node represents an immune parameter. Nodes are grouped by lineage and coloured by function, node size relates to number of relationships. (E) Plots depicting correlation between frequency of (left panel) IFN-γ+ γδ T cells and CD161-expressing γδ T cells; (middle panel) IFN-γ+ CXCL8-CD8 T cells and IFN-γ+ γδ T cells; and (right panel) CXCL8-producing CD4 and CD8 T cells. The data shown are a pool of longitudinal samples from 39 preterm babies where each circle represents an individual sample and on average there are 8 longitudinal samples/baby. Individual babies are in different colours.

Importantly, this trajectory was generally followed by all the babies, independent of the infant’s GA at birth. In fact, when we examined distance travelled along this trajectory, babies born at the earliest GA tended to exhibit a greater degree of movement over a similar time period to those born at later gestational ages (**Fig 2B**). tSNE analyses suggested that even though all babies progressed down this shared developmental pathway, samples taken from the same baby (represented by individual colours) appeared to retain unique features sufficient to cluster samples most closely according to their individual profile. This distinction of babies one from another was sustained throughout their time in the NICU (**Fig 2C**).

#### Associations between immune parameters

To identify any immune cell populations that may associate with each other (either positively or negatively) in our infant cohort, we performed Spearman correlation coefficients (R=>0.3 0r R=<-0.3) across our whole data set. We identified significant associations (p<0.008) between several immune parameters, many of which were to be expected (e.g. negative correlation between naïve and memory T cells). These associations were grouped into nodes based on cell type (see **Fig 2D**). Some populations (e.g. intermediate monocytes) showed no correlation with others and were subsequently lost from this network analysis. Examples of unanticipated positive correlations included the expression of CD161 on γδ T cells correlating with their production of IFN-γ [R=0.5] and between IFN-γ producing (CXCL8-) γδ and CD8 T cells [R=0.41] (**Fig 2E**). There was also a good correlation between CXCL8-producing CD4 and CD8 T cells [R=0.62]) (**Fig 2E**).

### 3.3 Potential Sources Controlling Immune Profile Variation

There were extensive differences identified between our preterm infants at birth and the equivalent adult immune profile. However, whilst some immune parameters exhibited rapid maturation over the time frame of our study, even in those infants born extremely preterm, others showed little change. Considering the links previously suggested between immune cell development, immune homeostasis and the microbiome ^20^, we considered the developing microbiome as a potential driver of these changes in immune profile. Similarly, many immune parameters are sexually dimorphic, at least in mice ^21^. Hence, we also considered the influence of sex on these developing immune profiles.

Microbiome analyses were performed on 10 stable infants (100% of eligible cohort); 10 unstable infants (63% of eligible cohort); and 11 unstable infants born to mothers with chorioamnionitis (85% of eligible cohort). Initially, we analysed the development of the microbiome by assessing the range of different taxa that appeared over time (longitudinal development of diversity). In the stable group, microbiome diversity increased at a rate of 0.56 units/week (95%CI 0.3-0.9). Conversely, there was significantly restricted growth in diversity in both the unstable (median 0.035 vs. 0.56 units/week, p<0.0001) and the unstable chorioamnionitis group (0.065 vs. 0.56, p<0.0001), with no significant difference in the rate of diversity progression between these cohorts (**Fig 3A**).

**Figure 3:**
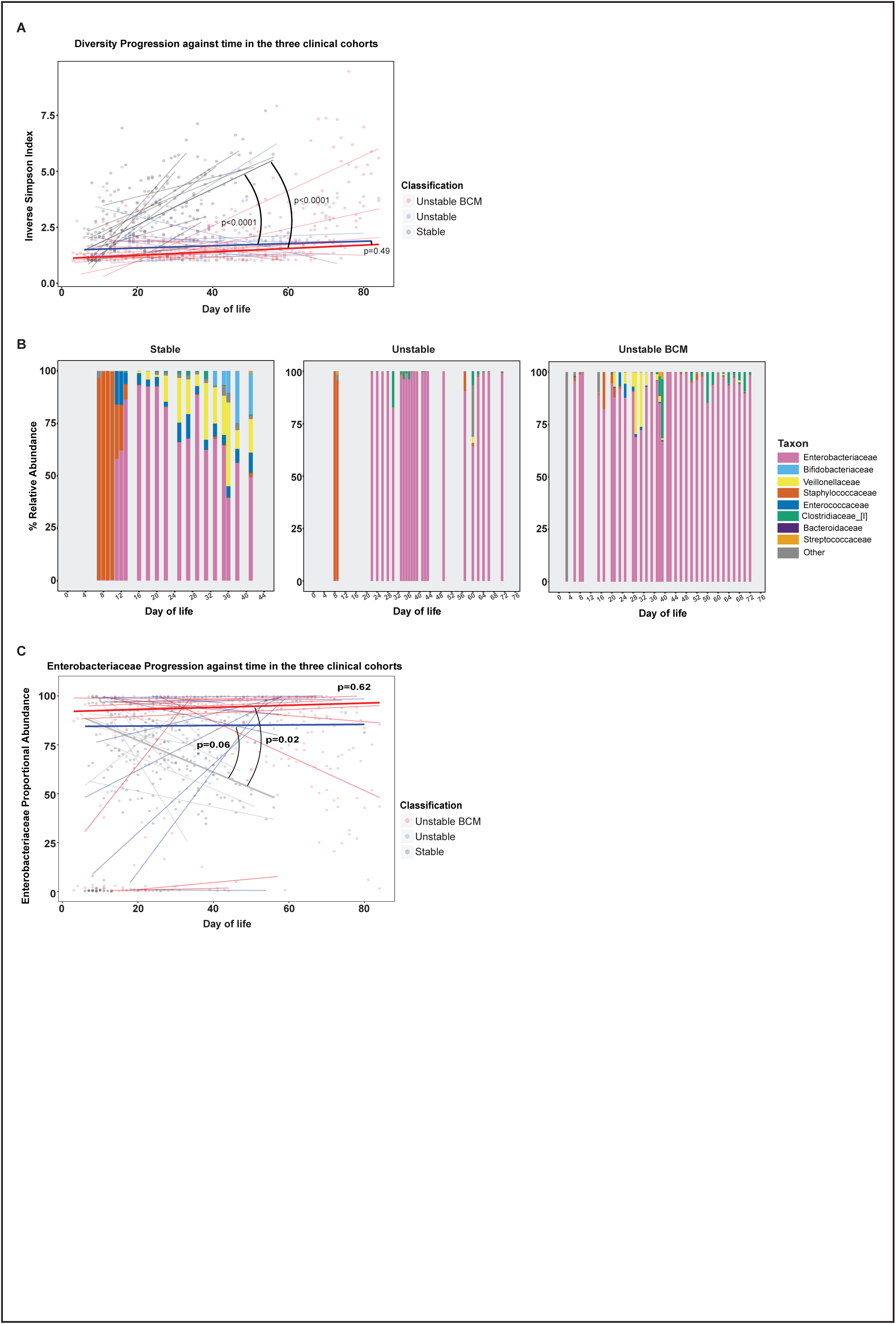
Microbiome development. Faecal samples (672) were longitudinally collected from 31 babies across the duration of the study. Bacterial DNA was extracted, and the 16S rRNA gene (hypervariable region V4) was amplified, sequenced and compared against the sequence database to ascertain the relative abundance of bacterial taxa in samples. Subjects were classified into three groups based on their clinical progress: stable, unstable, and unstable with chorioamnionitis. (A) The progression of diversity against postnatal age across the three groups (Stable – grey; Unstable – blue; Unstable with chorioamnionitis – red). Dot plot represents individual sample diversity indices. Thin lines represent individual subjects’ regression coefficients through their diversity data points, as assessed by Theil-Sen estimator. Thick lines represent the median/mean of the individual trajectories within each clinical group. (B) Stacked bar charts of relative abundance of taxa in typical subjects from the three clinical groups. Only taxa with >1% mean abundance across all samples represented. (C) *Enterobacteriaceae* relative abundance against postnatal age across the three groups (Stable – grey; Unstable – blue; Unstable with chorioamnionitis – red). Dot plot represents individual sample *Enterobacteriaceae* proportions. Thin lines represent individual subjects’ regression coefficients through their *Enterobacteriaceae* proportions data points, as assessed by Theil-Sen estimator. Thick lines represent the median/mean of the individual trajectories within each clinical group.

When assessing the emergence of taxa in our cohort, the family *Enterobacteriaceae* was the dominant family across all the groups. **Fig 3B** shows representative examples of taxa progression in individual babies from the three clinical classifications. When the longitudinal progression of *Enterobacteriaceae* was compared between the groups, *Enterobacteriaceae* relative abundance fell much quicker in the stable cohort (median 5.6%/week) than either the unstable (−5.6 vs. 0.16, p=0.02), or the unstable chorioamnionitis groups (−5.6 vs. 0.38 p=0.06). (**Fig 3C**).

As *Enterobacteriaceae* relative abundance fell in the stable cohort, a shift towards higher proportions of other families was observed. The next most prevalent were *Veillonellaceae* (which gradually increased over time) and *Bifidobacteriaceae*, (which increased from approximately day 30) having a maximum relative abundance of >1% in 9/10 and 10/10 subjects respectively (**Supplementary Fig 5**). Conversely, *Staphylococcaceae* were highly prevalent (>1% maximum abundance in 8/10 subjects) in early life, becoming almost absent beyond day 20. Other notable families (*Clostridiaceae* 8/10; *Enterococcaceae* 9/10), changed minimally over time, and generally maintained low median abundances. In the unstable groups, these consistent patterns of microbial community progression were not observed (**Supplementary Fig 5**), although a strikingly decreased prevalence of *Bifidobacteriaceae* was noted (38% of unstable subjects were never colonised with *Bifidobacteriaceae* i.e. never had a relative abundance in any sample >1%.).

When the microbiome structure was assessed in conjunction with the immune parameters, we could identify very few correlations between the two data sets (**Supplementary Fig 6**). The only weak association identified was between *Enterococcaceae* and B cells.

To try to establish additional factors driving the different immune profiles in our preterm infant cohort, we analysed for differences driven by sex. Perhaps surprisingly, the majority of immune parameters showed no significant sexual dimorphism, but there were three notable exceptions. Both CD31 and CD38 expression on CD4 T cells were significantly elevated in female infants compared to their male counterparts (**Supplementary Fig 7A/B**). In contrast, CD161 expression on CD8 T cells was higher in males and then subsequently decreased more dramatically in expression over time than in females (**Supplementary Fig 7C**).

#### 3.3.1 In-utero and post-natal exposure to Inflammation

Having shown limited associations between immune parameters and sex/microbiome we considered the effects of both pre- and post-natal exposure to infection on the developing immune system. Analyses of the whole immune profiles by PCA for the different clinical groups described in Table 2 identified distinct immune signatures in babies who had experienced different pre- and post-natal exposures to the stable infants (**Fig 4A/B**). Babies who had an unstable course or those born in the context of chorioamnionitis, showed elevated levels of the activation marker CD69 on all T cell populations (CD4, CD8 and γδ) as well as on NK cells (**Fig 4C-F**) when compared to babies who had a stable postnatal course. CD35, (also known to increase upon cell activation) was also elevated on both CD4 and CD8 T cells, significantly so in those infants exposed to chorioamnionitis (**Supplementary Fig 8A**). There was no difference in CD25 expression, however, on either CD4 or CD8 T cells between the different clinical groups.

**Figure 4:**
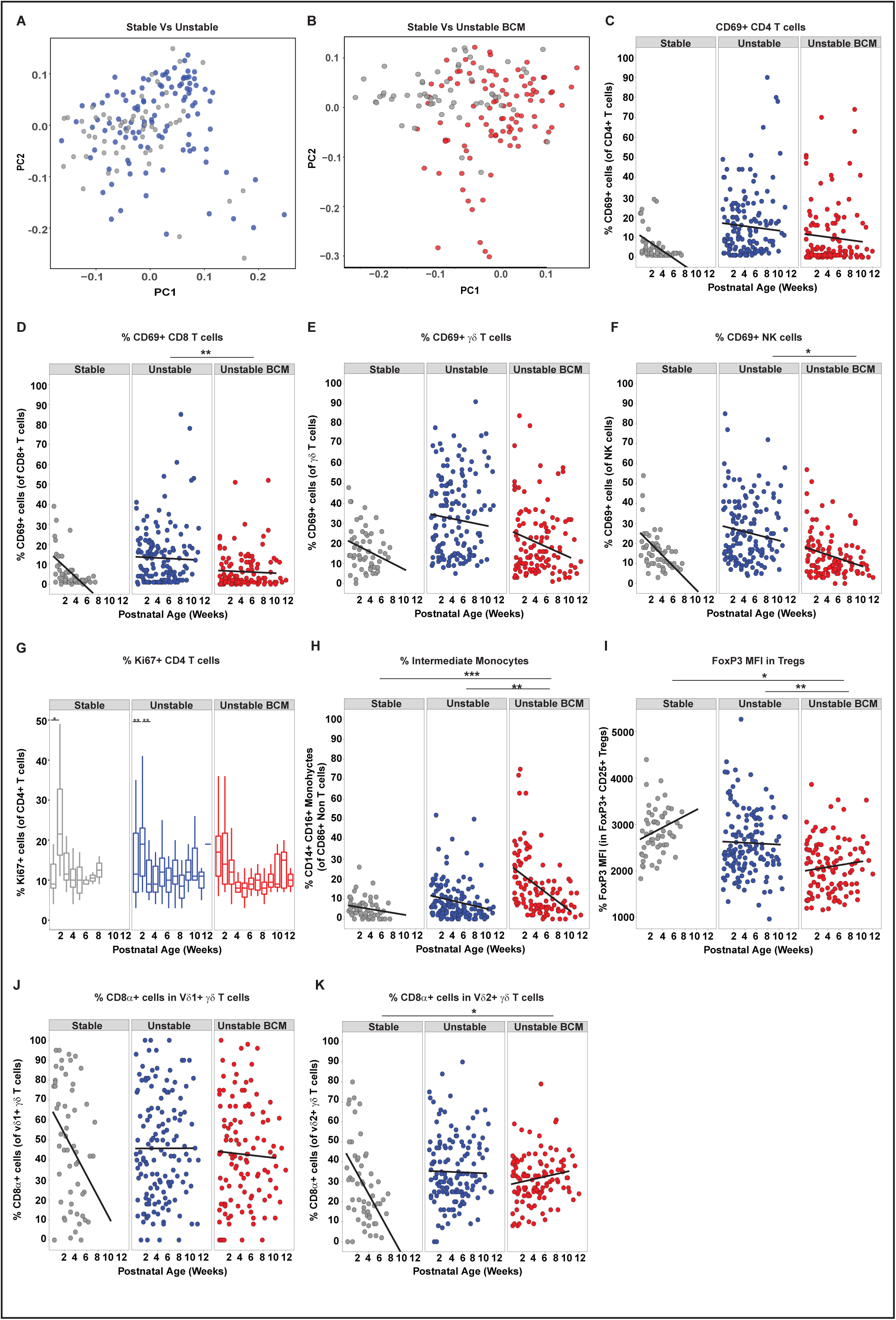
Immune parameters altered by different pre- and/or post-natal exposures. Longitudinal PBMC samples from 39 preterm babies were phenotyped for 186 different immune populations by flow cytometry following surface and intracellular staining. For cytokine detection, samples were activated in vitro with PI (4 h, in the presence of BFA) prior to staining. PCA of 186 immune parameters from longitudinal samples derived from: (A) Stable (grey circles; n=10 babies) and Unstable (blue circles, n=16) babies (B) Stable (grey circles; n=10) and Unstable BCM (red circles, n=13) babies. In Figures C-F, scatter plots depict frequencies of CD69 positive cells in Stable babies (left panel; grey circles, n=10), Unstable babies (middle panel; blue circles, n=16) and Unstable BCM babies (right panel; red circles, n=13) as a function of postnatal age: (C) CD69+ CD4 T cells, (D) CD69+ CD8 T cells, (E) CD69+ γδ T cells, (F) CD69+ NK cells are depicted from the three clinical groups. (G) Proliferation (as depicted by the boxplot showing frequency of Ki67 expression in CD4+ T cells) was elevated immediately post birth in infants born to mothers with chorioamnionitis whereas stable (and to a lesser extent unstable) infants showed a proliferative burst at around 14 days of age. Compared to Stable babies, Unstable and Unstable BCM babies show significantly higher frequencies of Intermediate monocytes (H) and significantly lower FoxP3 MFI in Tregs (I) and higher frequencies of CD8α expressing γδ T cells over time in both (J) vδ1+ γδ T cells, (K) vδ2+ γδ T cells. Data shown are a pool of longitudinal samples from each clinical group where on average there are 8 samples/baby in the Stable cohort (n=10) and 9 samples per baby in the Unstable cohort (n=16) and the Unstable BCM cohort (n=13). For all figures except (G) *** p< 0.001, **p < 0.01 and *p< 0.05 as determined by linear mixed effect modelling using the lmer package in R. For (G) **p < 0.01 and *p< 0.05 as determined by a non-parametric Wilcoxon matched-pairs signed rank test.

Stable infants exhibited low levels of proliferation at birth (<10% of CD4 T cells were Ki67 positive). However, there was a consistent proliferative burst (to nearly a quarter of CD4 T cells) around 2 weeks after birth which rapidly returned to post birth levels, seen in nearly all infants (**Fig 4G**). Elevated CD4 T cell proliferation was also observed immediately post birth in infants born to mothers with chorioamnionitis (**Fig 4G**). Similarly, elevated levels of intermediate monocytes, particularly in the immediate perinatal period, were seen in babies born to mothers with chorioamnionitis (p=0.0004, **Fig 4H**) both suggestive of in-utero activation. However, levels of HLADR expression on these perinatal monocytes were significantly lower than that seen in stable infants (**Supplementary Fig 8B**). Despite no significant changes in regulatory T cell (T-reg) numbers between the different clinical groups (data not shown), FOXP3 expression was significantly reduced on T-regs isolated from infants born to mothers with chorioamnionitis (**Fig 4I**). CD8 expression was maintained on γδ T cells (both Vδ2 and Vδ1) in unstable infants compared to those with a stable clinical course (where levels decreased rapidly post birth, **Fig 4J,K**).

To identify any changes in functionality between the different clinical groups, cytokine production following *in vitro* stimulation with PMA and Ionomycin was assessed. The ability of CD4 T cells to produce TNF-α and IL-2 (**Fig 5A/B**) upon *in vitro* stimulation was indistinguishable between the three clinical groups yet, in contrast, the ability to produce CXCL8 was severely and significantly reduced in both the chorioamnionitis and unstable groups when compared to stable infants (**Fig 5C**). It has been previously suggested that the capacity of T cells to produce TNF-α and CXCL8 was reciprocally regulated based on GA ^22^. However, we did not see significant changes in the capacity of CD4 T cells to make TNF-α or CXCL8 based on their GA at birth in this cohort (**Supplementary Fig 9A/B**). Similarly, babies born at the same GA exhibited very different percentages of T cells with the capability to produce either CXCL8 or TNF-α (**Supplementary Fig 9C/D**). In every case, despite an equivalent GA at birth, those babies exhibiting the lowest levels of CXCL8-producing T cells (but not TNF-α producing T cells) were those infants with an unstable clinical course (**Supplementary Fig 9C/D**). Furthermore, those infants demonstrating the lowest levels of CXCL8-producing T cells at birth were those that subsequently developed the most severe infections despite an equivalent capacity to produce TNF-α or IL-2 (**Fig 5D-F**). When compared to stable infants, significantly lower CXCL8-producing CD4 T cells were seen in infants with clinically suspected sepsis ^3^ (n=16, p=0.007) and in those infants with microbiologically confirmed sepsis caused by either coagulase negative Staphylococcus (CONS, n=9, p=0.039) or other bacteria (n=4, p=0.0007) such as Group B streptococcus [GBS], *Escherichia* coli or *Streptococcus anginosus*.

**Figure 5:**
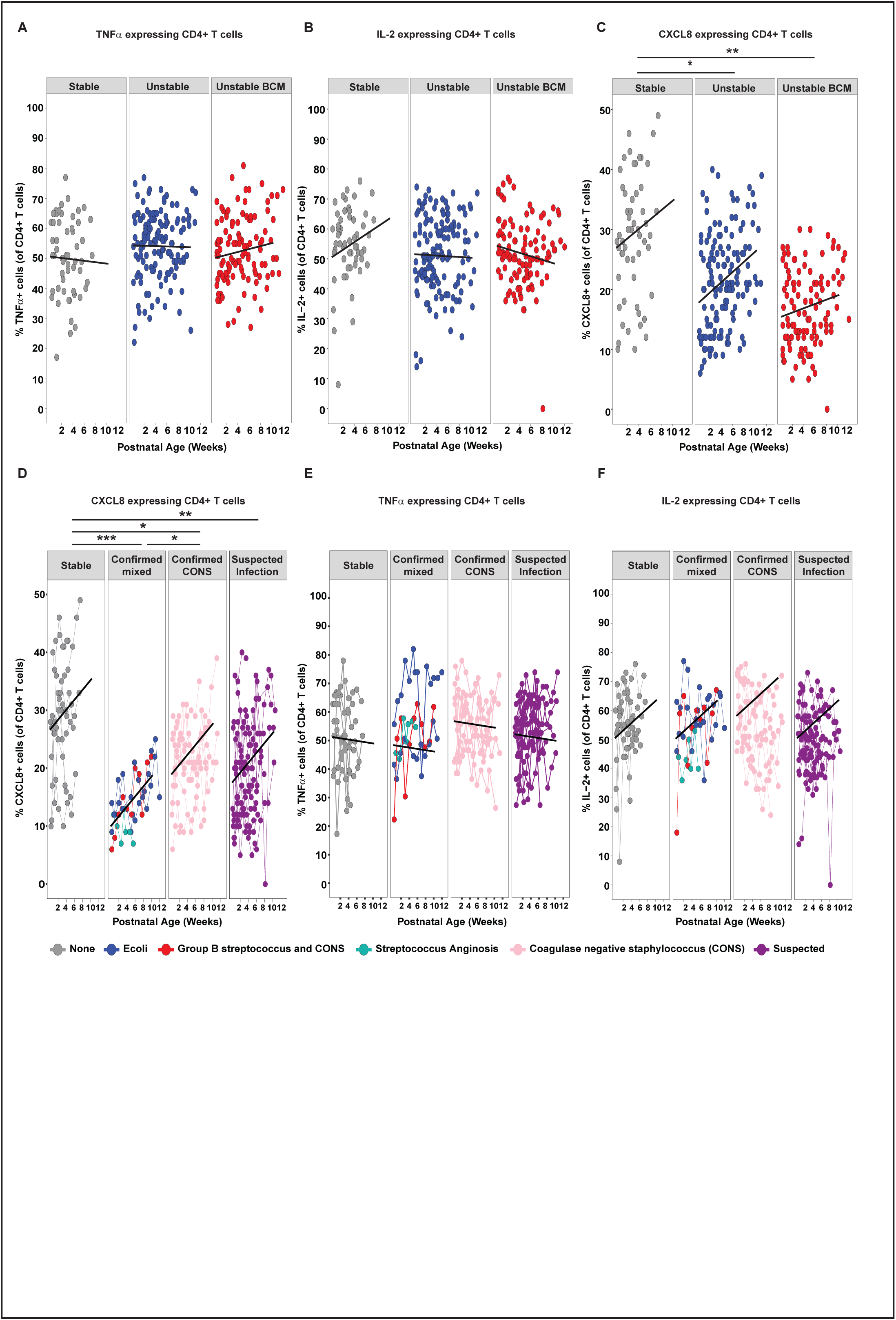
T cells from unstable infants have a significantly reduced ability to produce CXCL8 compared to stable babies. Longitudinal PBMC samples from 39 preterm babies were activated in vitro with PI (4 h, in the presence of BFA) and expression of CXCL8, TNF-α and IL-2 assessed by flow cytometry. Compared to Stable babies, Unstable and Unstable BCM babies show a significantly reduced ability to produce CXCL8 but not TNF-α or IL-2 as depicted by scatter plots showing frequencies of (A) TNF-α, (B) IL-2 and (C) CXCL8 in Stable babies (left panel; grey circles, n=10), Unstable babies (middle panel; blue circles, n=16) and Unstable BCM babies (right panel; red circles, n=13) as a function of postnatal age. In Figures A-C, data shown are a pool of longitudinal samples from each clinical group where each circle represents a longitudinal sample from an individual baby and on average there are 6 samples/baby in the Stable cohort and 9 samples per baby in the Unstable cohort and the Unstable BCM cohort. Preterm babies with the lowest levels of CXCL8 developed the most severe infections as depicted by scatter plots showing the frequencies of (D) CXCL8, (E) TNF-α and (F) IL-2 in Stable babies (far left panel; grey circles, n=10 babies with mean of 8 longitudinal samples/baby), babies with microbiologically confirmed serious infections (second panel; n=4 with mean of 11 longitudinal samples/baby), babies with coagulase negative staphylococcus infections (CONS) (third panel, n=9 with mean of 10 longitudinal samples/baby) and babies with suspected infections (far right panel; n=16 with mean of 7 longitudinal samples/baby). The colour of the circles depicts the type of infection as described in the figure. In figures D-F, each circle represents a longitudinal sample from an individual baby and linked circles represent longitudinal samples from the same baby. *** p< 0.001, **p < 0.01 and *p< 0.05 as determined by linear mixed effect modelling using the lmer package in R

## 4 Discussion

The consequences of preterm birth on immune development and function are not well established ^23^. In this study, we demonstrate that preterm babies, even those born at 23 and 24 weeks’ gestation, were capable of rapidly acquiring several components of immune function. The immune profiles for babies born at earlier gestation changed more dramatically over a similar time period to those born at later gestations which suggests that extremely preterm babies are capable of rapid progression to ‘catch up’ immune function. This corroborates data suggesting that the immune profiles of preterm and term babies converge in a similar time frame ^4^ and indeed that rapid immune development in term babies follows a set trajectory^5^. However, despite this immune development, some babies developed infection. Strikingly, this was associated with a reduced T cell capacity to produce CXCL8, highlighting the CXCL8 pathway as a potential therapeutic target for the treatment of infection in preterm babies.

Immune development occurred apparently independently of changes within the developing microbiome. Prolonged broad-spectrum antibiotic treatment of the unstable infants likely transformed the developing microbiome and yet those infants were still capable of immune maturation along the established trajectory. Given the generally naïve nature of the neonatal immune system, it is conceivable that exposure to *any* environmental stimulus is sufficient to drive the observed maturation of the immune system during this period. Indeed, the sudden increase in proliferation of CD4 T cells and other T cells, that we observed at around 2 weeks of age may be related to microbial colonisation. Rapid colonisation with myriad microbial species would most likely far eclipse any differences driven by specific individual colonisers. Alternatively, microbial-immune relationships may exist below family level. Similarly, although the stable infants showed some consistency in microbial profiles with decreased *Enterobacteriacae* over time, these were replaced by a variety of different obligate anaerobes.

Whilst mode of birth has been associated with the development of specific microbiomes ^24^, we were unable to dissociate mode of birth from other exposures as all stable (but not all unstable) infants in our cohort were born by caesarean section. Similarly, whilst breast-feeding (either wholly or partially) is also known to significantly influence microbiome development ^25^, almost all of our babies received at least some maternal breast milk.

Aspects of both innate and adaptive immunity developed rapidly during the period of study. B cells increased in numbers after birth as did γδ T cells whose function (as determined by IFN-γ production) was enhanced over time. We identified a decrease in CD8 expression on γδ T cells, but interestingly, only in stable babies. This suggests either clonal expansion of an existing CD8+ population and/or increased CD8 expression upon activation in those infants experiencing episodes of infection. It is intriguing that we see significant changes in the γδ T cell population in our cohort as we and others have suggested these cells are of particular importance in providing protection to infants in early life ^26 27 28^. In contrast, αβ CD4 T cells showed scant, if any, development over this time period. There was little change in their ability to produce CXCL8, which, if anything, increased over the time frame, presumably as a result of increased T cell export from the thymus post birth ^29 30^ and they were still unable to produce IFN-γ upon ex vivo stimulation. This relative inability to produce IFN-γ may be related to hypermethylation of the IFN-γ promoter previously observed in human newborn infants ^31^. Although, we did not see any significant increase in IFN-γ producing CD8 T cells over postnatal age there was a significant association between IFN-γ producing γδ T cells and IFN-γ producing CD8 T cells suggesting some maturation in the latter, albeit at a slower rate than that observed for γδ T cells.

In this study, an individual baby’s subsequent clinical course could not have been predicted at the time of birth. Babies with an unstable postnatal clinical course and those born in the context of chorioamnionitis both succumbed to higher rates of microbiologically confirmed sepsis and chronic lung disease when compared to the stable cohort. Our data highlight the influence of both the in-utero environment and the post-natal clinical course on the developing immune profiles of preterm infants. Bacteria implicated in chorioamnionitis (such as *E. coli*) have also been shown to produce histone modifying enzymes that down regulate protein expression with increased evidence that sepsis can induce epigenetic changes that alter immune responses through a range of mechanisms ^32,33^.

An activated immune profile at birth, as suggested by several parameters (for example expression of CD69, CD35 and proliferation) and previously observed ^34^ was seen in infants born to mothers with chorioamnionitis. Furthermore, significantly elevated intermediate monocytes were also observed in this group. However, although increased in number, their functional potentials may be poor as we identified significantly reduced HLADR expression on these cells, as has also been observed previously ^35^. Indeed, monocytes developing in the context of chorioamnionitis have shown hyporesponsiveness to different stimuli ^35,36^ perhaps explaining the increased susceptibility to early onset sepsis observed in this patient group ^37^. Similarly, those infants born to mothers with chorioamnionitis also exhibited significantly lower levels of FOXP3 expression in T-regs suggesting reduced functionality ^38^.

Sex is known to have a significant bearing on neonatal outcomes with male preterm infants experiencing more adverse outcomes than females ^39^, and yet we saw very few significant differences in immune parameters between male and female infants. The few immune differences that were identified may, therefore, be functionally relevant. Both CD31, and its binding partner, CD38, were significantly lower in male CD4 T cells compared to their female counterparts, suggesting the whole pathway may be less efficient. Interestingly, CD38 has been shown to enhance T cell (and NK) activation by contributing to immune synapse formation ^40 41^ and CD31 deficient mice show enhanced susceptibility to LPS induced endotoxic shock ^42^. The only other factor that appeared to be sexually dimorphic was CD161 expression on CD8 T cells, which decreased in both male and female infants postnatally (albeit from an increased starting level in males). The expression of CD161 is thought to evoke a particular functional profile ^43^ irrelevant of the cell type. It is interesting therefore, that expression was enhanced postnatally on NK cells and γδ T cells (where it correlated with IFN-γ production) but decreased on CD8 T cells. It is possible that some CD161+ CD8+ T cells represent mucosal associated invariant T (MAIT) cells and thus the rapid decrease may be associated with migration out of the blood into the intestine.

The principal effector chemokine of neonatal CD4 T cells is CXCL8 ^10^. CXCL8 expression was highly divergent between infants and adults, and there was a strong correlation between CXCL8 producing CD4 and CD8 T cells in our babies. Levels of CXCL8-producing αβ T cells did not decrease over the time period studied towards the adult levels, despite reductions in CXCL8-producing NK and γδ T cells. This notwithstanding, when compared to stable infants, both groups of unstable infants exhibited significantly reduced CXCL8-producing CD4 and CD8 T cells. There may be several reasons for this observation. It is possible that these cells are somehow exhausted in the unstable cohorts. This is unlikely to be the case as the ability of the same cells to produce TNF-α or IL2 was not reduced. Furthermore, both infection and inflammation may themselves be implicated. In this study, we show that babies born in the context of chorioamnionitis (inflammation) and those with repeated episodes of infection have lower CXCL8 responses but we do not know what mechanisms link this association. The observation that CXCL8 and TNF-α production may be reciprocally associated with GA ^22^ and that unstable infants in this cohort were generally born at an earlier GA, may also be an explanation. However, amongst this cohort, there appeared to be no correlation between GA and the ability of CD4 T cells to produce CXCL8 or TNF-α around birth. Nevertheless, a reduced number of CXCL8-producing T cells was consistently observed in those infants with an unstable clinical course who were more likely to develop serious postnatal complications. This inability to mount adequate T cell CXCL8 responses at birth (for whatever reason) may therefore predict a poor outcome. This finding may have particular clinical relevance, both identifying CXCL8 as a potential therapeutic target but also as a biomarker to predict subsequent outcomes for babies born prematurely, heretofore a challenging area.

## 7. Supplementary figure legends

**Supplementary Figure 1: Schematic of study design**

**Supplementary Figure 2: Comparison of neonatal and adult immune parameters**

Longitudinal PBMC samples from 39 preterm babies were phenotyped for 186 different immune populations by flow cytometry following surface and intracellular staining. For cytokine detection, samples were activated in vitro with PI (4 h, in the presence of BFA) prior to staining. (A) Bar chart depicting the deviation of the neonatal immune profile from adult levels. Plot represents Log2 fold difference of mean neonatal immune cell subset frequencies (n=39 infants) relative to mean adult levels (n=9 adults). Colour indicates –Log10 adjusted p value of difference in mean values between the two groups. Populations highlighted in blue text indicate adaptive CXCL8+ cells and those highlighted in red indicate innate (NK and γδ) CXCL8+ cells.

**Supplementary Figure 3: Immune parameters that increase with age**

The mean fluorescence intensity of HLADR expression in classical monocytes and the frequencies of CD19+ B cells, NK cells and NKG2D+ NK cells were examined in longitudinal PBMC samples from 39 preterm babies by flow cytometry. (A) Scatter plot depicting actual numbers/ml of CD19+ B cells. (B) Representative histograms depicting HLADR MFI in classical monocytes with increasing postnatal age in a Stable (left panel), Unstable (middle panel) and Unstable BCM (right panel) baby. Scatter plots depicting actual numbers/ml of (C) NK cells and (D) NKG2D+ NK cells in preterm babies as a function of postnatal age. Data shown in Figures (A) and (B-C) are a pool of longitudinal samples from 39 preterm babies where each circle represents an individual sample. *** p< 0.001, **p < 0.01 and *p< 0.05 as determined by linear mixed effect modelling using the lmer package in R

**Supplementary Figure 4: IFN-γ expression in γδ and CD4 T cells**

Longitudinal PBMC samples from 39 preterm babies were activated in vitro with PI (4 h, in the presence of BFA) and expression of IFN-γ in γδ or CD4 T cells assessed by flow cytometry. Dot plots showing representative staining of IFN-γ expression by (A) γδ T cells and (B) CD4+ T cells in longitudinal samples from a single preterm baby. The postnatal age of the baby at the time the sample was taken is indicated above each dot plot.

**Supplementary Figure 5: Microbiome development**

Faecal samples (672) were longitudinally collected from 31 babies across the duration of the study. Bacterial DNA was extracted, and the 16S rRNA gene (hypervariable region V4) was amplified, sequenced and compared against the sequence database to ascertain the relative abundance of bacterial taxa in samples. We have utilised our own curated database, created to focus upon species previously encountered in preterm intestinal microbiome studies. Mean abundance of minority taxa was plotted against time for all samples and subjects. Dot plot represents individual samples - colours represent samples originating from same subject; subjects who were never colonised with that taxon (i.e. relative abundance always <1%) are uniformly coloured in black.

**Supplementary Figure 6: Immune microbiome correlations**

Heat map depicting aggregate intra-individual spearman correlations between microbiome parameters and immune parameters. Samples from 24 babies were included in this analysis to ensure > 4 samples were included per baby.

**Supplementary Figure 7: Sexual dimorphism in immune parameters**

Surface expression of CD31, CD38 on CD4 T cells and CD161 on CD8 T cells was examined by flow cytometry in longitudinal PBMC samples from 39 preterm babies. Compared to males, female babies express significantly higher levels of CD31 and CD38 on CD4+ T cells and significantly lower levels of CD161 on CD8+ T cells as depicted by scatter plots showing frequencies of (A) CD31 and (B) CD38 in CD4 T cells and (C) CD161 on CD8+ T cells in males (left panel, n=23 babies with mean of 8 samples/baby) compared to females (right panel, n=16 with mean of 8 samples/baby) as a function of postnatal age. Data shown are a pool of longitudinal samples from 39 preterm babies where each circle represents an individual sample. *** p< 0.001, **p < 0.01 and *p< 0.05 as determined by linear mixed effect modelling using the lmer package in R.

**Supplementary Figure 8: CD35 expression in CD4 and CD8 T cells and HLADR MFI in intermediate monocytes**

The frequency of CD35-expressing CD4 and CD8 T cells and HLADR MFI in monocytes was examined by flow cytometry in longitudinal PBMC samples from 39 preterm babies. Scatter plots depicting the frequency/MFI (as indicated) of (A) CD35+ CD4+ T cells, (B) CD35+ CD8+ T cells and (C) HLADR MFI in intermediate monocytes in Stable babies (left panel; grey circles, n=10), Unstable babies (middle panel; blue circles, n=16) and Unstable BCM babies (right panel; red circles, n=13) as a function of postnatal age. Data shown are a pool of longitudinal samples from 39 preterm babies where each circle represents an individual sample and on average there are 8 longitudinal samples/baby. *** p< 0.001, **p < 0.01 and *p< 0.05 as determined by linear mixed effect modelling using the lmer package in R.

**Supplementary Figure 9: T cell expression of CXCL8 or TNF-α is not dependent on gestational age at birth**

Longitudinal PBMC samples from 39 preterm babies were activated in vitro with PI (4 h, in the presence of BFA) and expression of CXCL8 and TNF-α in CD4+ T cells was assessed by flow cytometry. Scatter plots show frequencies of CD4 T cells expressing (A) TNF-α and (B) CXCL8 in preterm babies as a function of corrected gestational age (cGA). Data shown are a pool of samples taken in the first week of life from 39 preterm babies where each circle represents an individual sample. Linked samples indicate consecutive samples from the same baby within the first week of birth. Circles are coloured based on the gestational age of the baby at birth. The colour gradient is from dark (babies born at earlier gestational ages) to light (babies born at later gestational ages). Unstable and Unstable BCM babies born at the same gestational age as stable babies consistently express lower frequencies of CXCL8 but not TNF-α as depicted by scatter plots showing frequencies of CD4+ T cells expressing: (C) CXCL8 and (D) TNF-α as a function of cGA in babies born at 29 weeks cGA (left panel), 30 weeks cGA (middle panel) and 31 weeks cGA (right panel). In figures C-D, circles depict individual samples from each baby. Linked samples are longitudinal samples from the same baby. Grey circles represents Stable babies, blue circles represent Unstable babies and Red circles represent Unstable BCM babies. *** p< 0.001, **p < 0.01 and *p< 0.05 as determined by linear mixed effect modelling using the lmer package in R.

**Supplementary Table 1.**

| Marker | Fluorochrome | Company | Catalogue No. | Clone | Dilution |
| --- | --- | --- | --- | --- | --- |
| **Surface markers** |  |  |  |  |  |
| Human<br>Trustain Fcα | N/A | Biologend | 422302 | N/A | 2.5ul/test |
| CD3 | AF700 | Biologend | 317339 | OKT3 | 1:200 |
| CD4 | PE-Cy7 | Biologend | 317413 | OKt4 | 1:50 |
| CD4 | BV786 | Biologend | 317442 | Okt4 | 1:50 |
| CD8 | BV510 | Biologend | 301047 | RPA-T8 | 1:400 |
| Vδ1 | FITC | Thermo Scientific | TCR2730 | TS8.2 | 1:100 |
| Vδ2 | PerCP | Biologend | 331410 | B6 | 1:50 |
| CD45RA | BV786 | Biologend | 304139 | HI 100 | 1:50 |
| CD45RA | BV650 | Biologend | 304135 | H100 | 1:250 |
| CCR7 | BV650 | Biologend | 353233 | GO43H7 | 1:50 |
| CD95 | BV421 | Biologend | 305623 | dx2 | 1:50 |
| CD25 | PE | Biologend | 356103 | MA-251 | 1:50 |
| Ki67 | BV421 | Biologend | 350505 | Ki-67 | 1:50 |
| CD27 | PE | Biologend | 356103 | MA-251 | 1:50 |
| CD19 | BV786 | Biologend | 302239 | HIB 19 | 1:100 |
| CD14 | BV421 | Biologend | 301829 | M5E2 | 1:100 |
| CD16 | APC/CY7 | Biologend | 301819 | M5E2 | 1:100 |
| CD14 | APC/CY7 | Biologend | 302017 | 3g8 | 1:100 |
| CD16 | PE-Cy7 | Biologend | 302015 | 3g8 | 1:200 |
| HLADR | AF488 | Biologend | 307619 | L243 | 1:100 |
| CD86 | PE | Biologend | 305405 | IT2.2 | 1:100 |
| CD40 | BV510 | Biologend | 334329 | 5c3 | 1:100 |
| CD1c(BDCA-1) | APC | Biologend | 331523 | L161 | 1:100 |
| CD56 | BV650 | Biologend | 362532 | 5.1h11 | 1:50 |
| CD123 | PerCP Cy5.5 | Biologend | 306015 | 6H6 | 1:50 |
| CD303 | FITC | Biologend | 307619 | L243 | 1:100 |
| CD11c | BV650 | Biologend | 301637 | 3.9 | 1:50 |
| Pan $\gamma\delta$ | PE-Cy7 | Biologend | 331222 | B1 | 1:50 |
| NKG2D | APC | Biologend | 320807 | 1D11 | 1:50 |
| CD56 | BV650 | Biologend | 362532 | 5.1h11 | 1:50 |
| CD56 | BV421 | Biologend | 362551 | 5.1h11 | 1:200 |
| CD69 | PerCP-Cy5.5 | Biologend | 310925 | FN50 | 1:50 |
| CD161 | AF488 | Biologend | 339923 | HP-3G10 | 1:50 |
| CD161 | BV510 | Biologend | 301047 | HP-3G10 | 1:40 |
| CD38 | BV650 | Biologend | 356619 | HB-7 | 1:100 |
| CD31 | PerCP-Cy5.5 | Biolegend | 303131 | WM59 | 1:400 |
| CD35 | AF647 | BD | 565329 | E11 | 1:50 |
| Intracellular markers: cytokines |  |  |  |  |  |
| IFNγ | BV421 | Biolegend | 502531 | 4S.B3 | 1:50 |
| TNFα | PerCP-Cy5.5 | Biolegend | 502925 | MAB11 | 1:50 |
| IL17A | AF647 | Biolegend | 512309 | BL168 | 1:50 |
| IL-13 | PE | Biolegend | 501903 | JES10-5A2 | 1:50 |
| IL-8 | FITC | Biolegend | 511406 | E8N1 | 1:50 |
| IL-2 | PE-CY7 | Biolegend | 500325 | MQ1-17H12 | 1:50 |
| IL-4 | PE | Biolegend | 500808 | MP425d2 | 1:50 |
| IL-10 | BV421 | Biolegend | 501421 | Jes3-97d | 1:50 |
| Intracellular markers: transcription factors |  |  |  |  |  |
| FoxP3 | APC | Biolegend | 320214 | 259 d | 1:50 |

