## Supplementary figures and images for "Perinatal Inflammation Influences but Does Not Arrest Rapid Immune Development in Preterm Babies"

### All Supplementary data

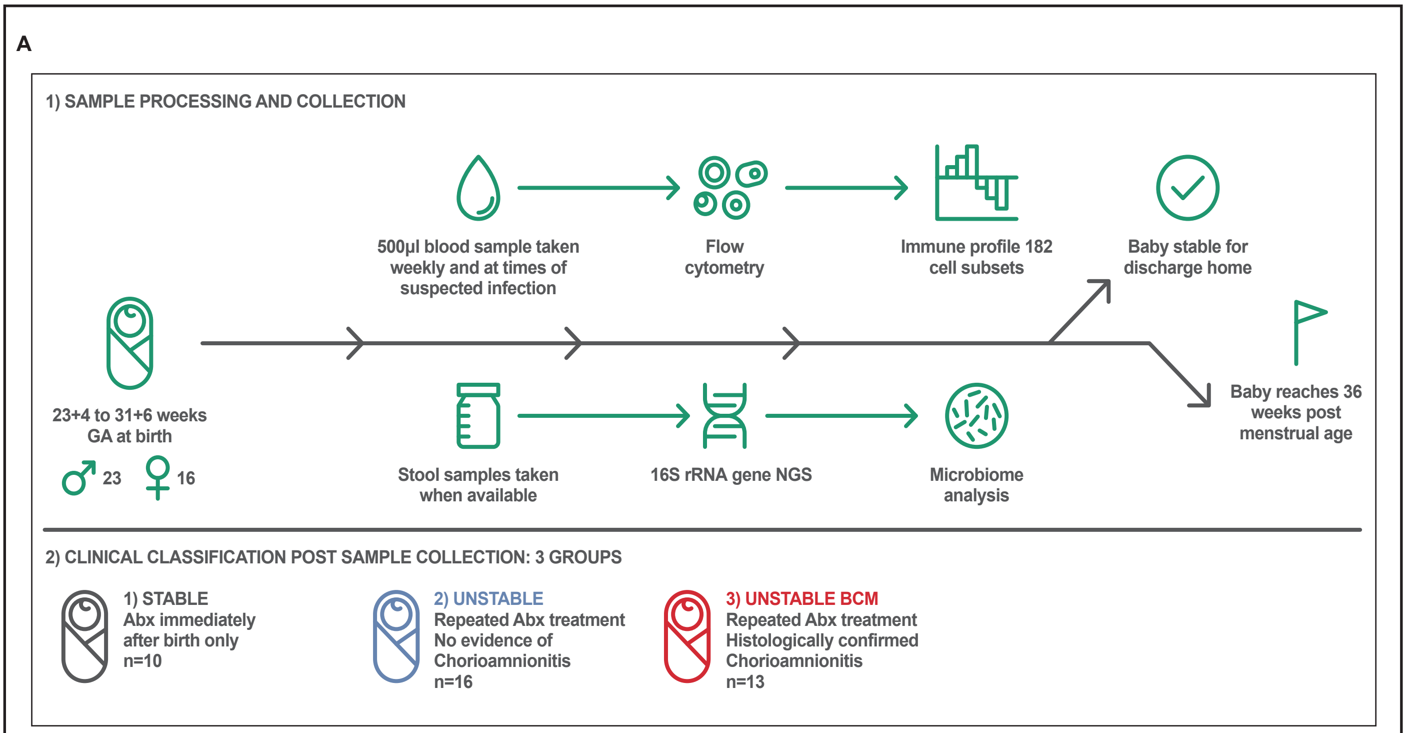

**Supplemnetary Figure 2**

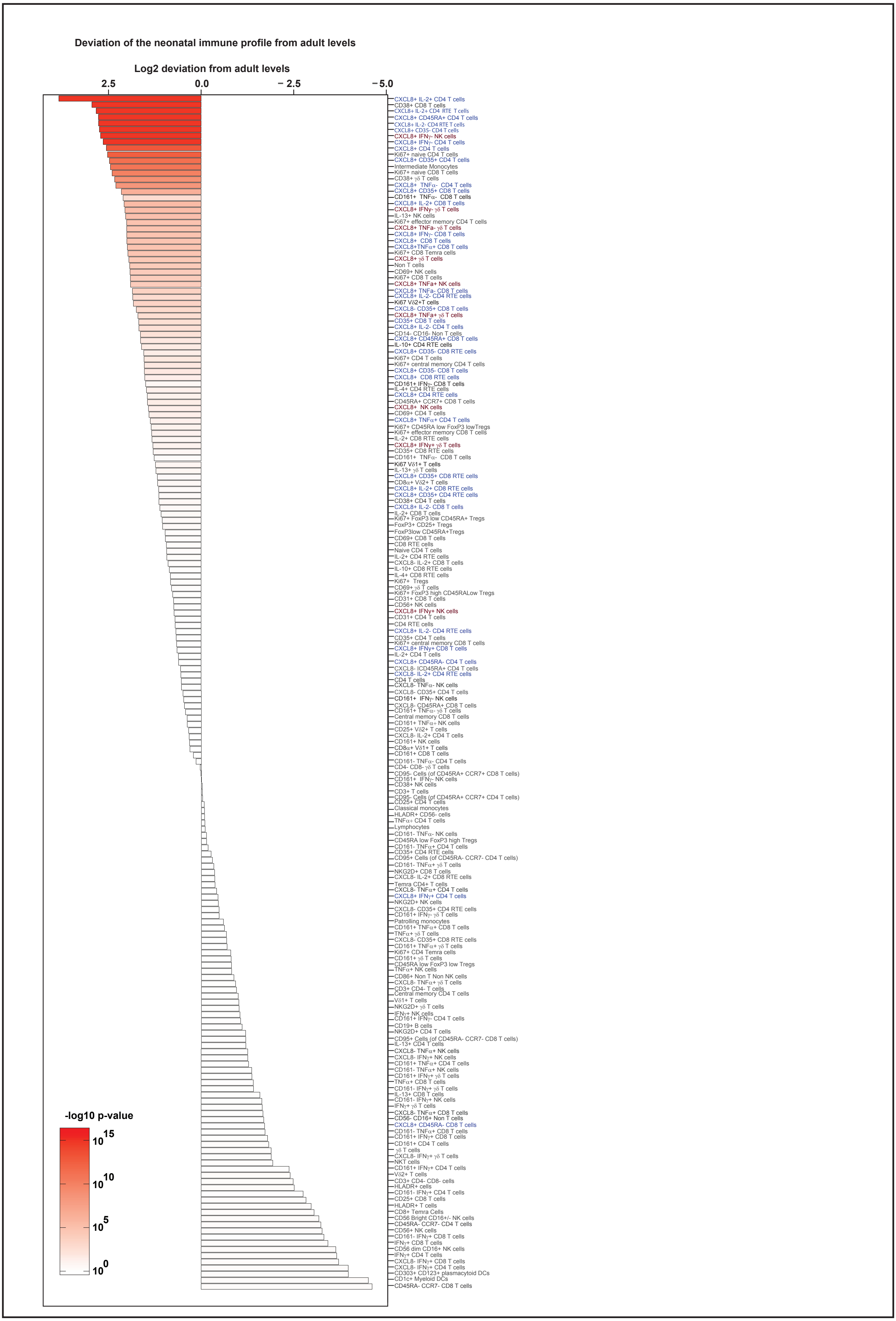

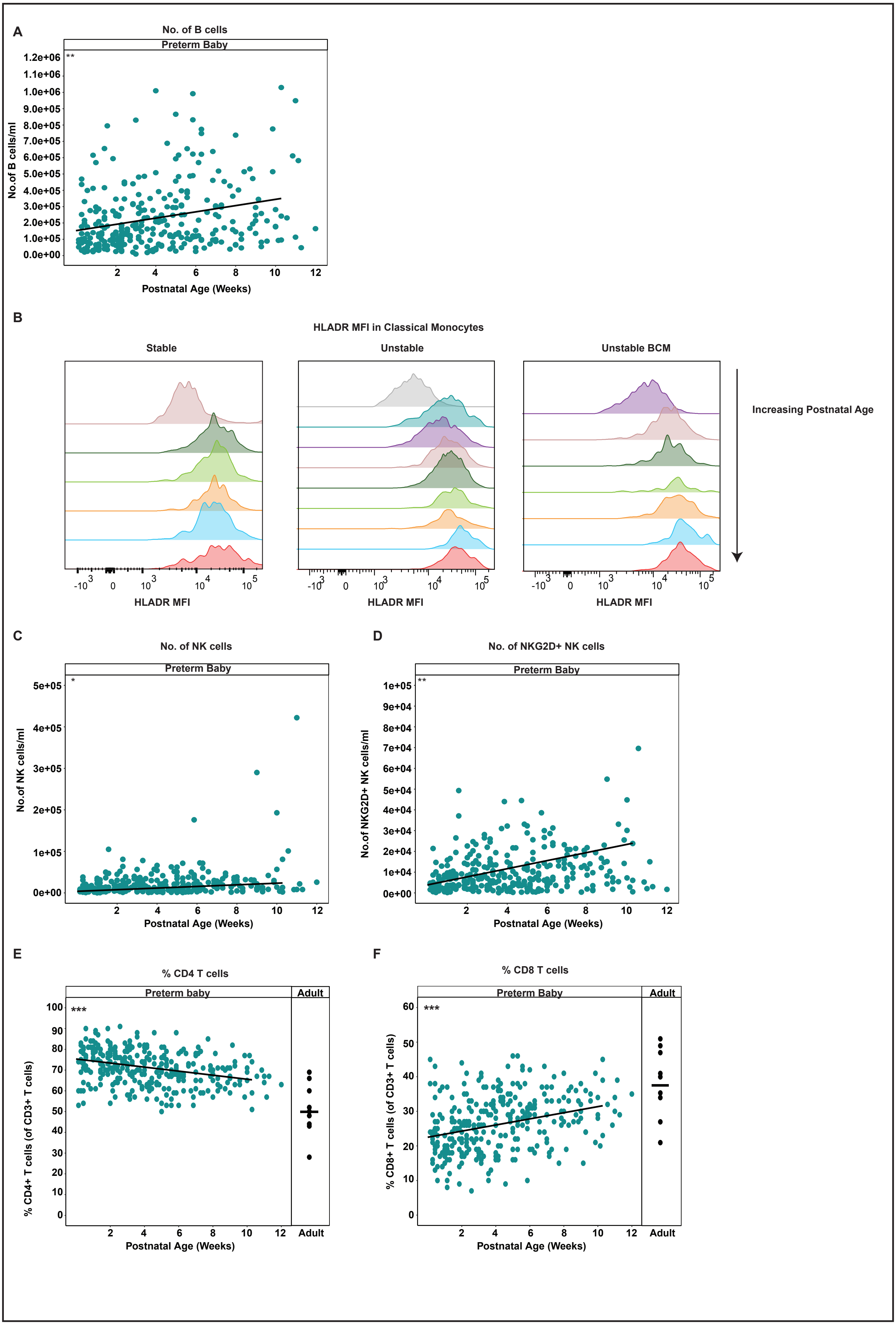

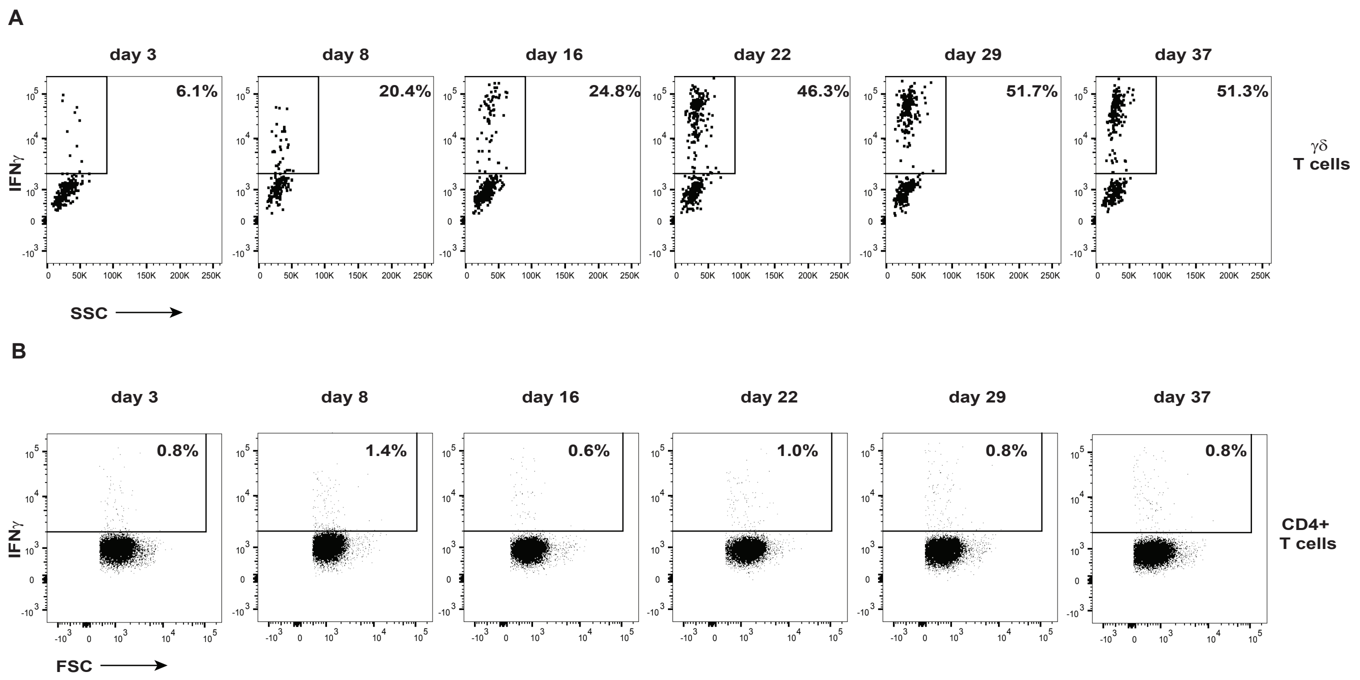

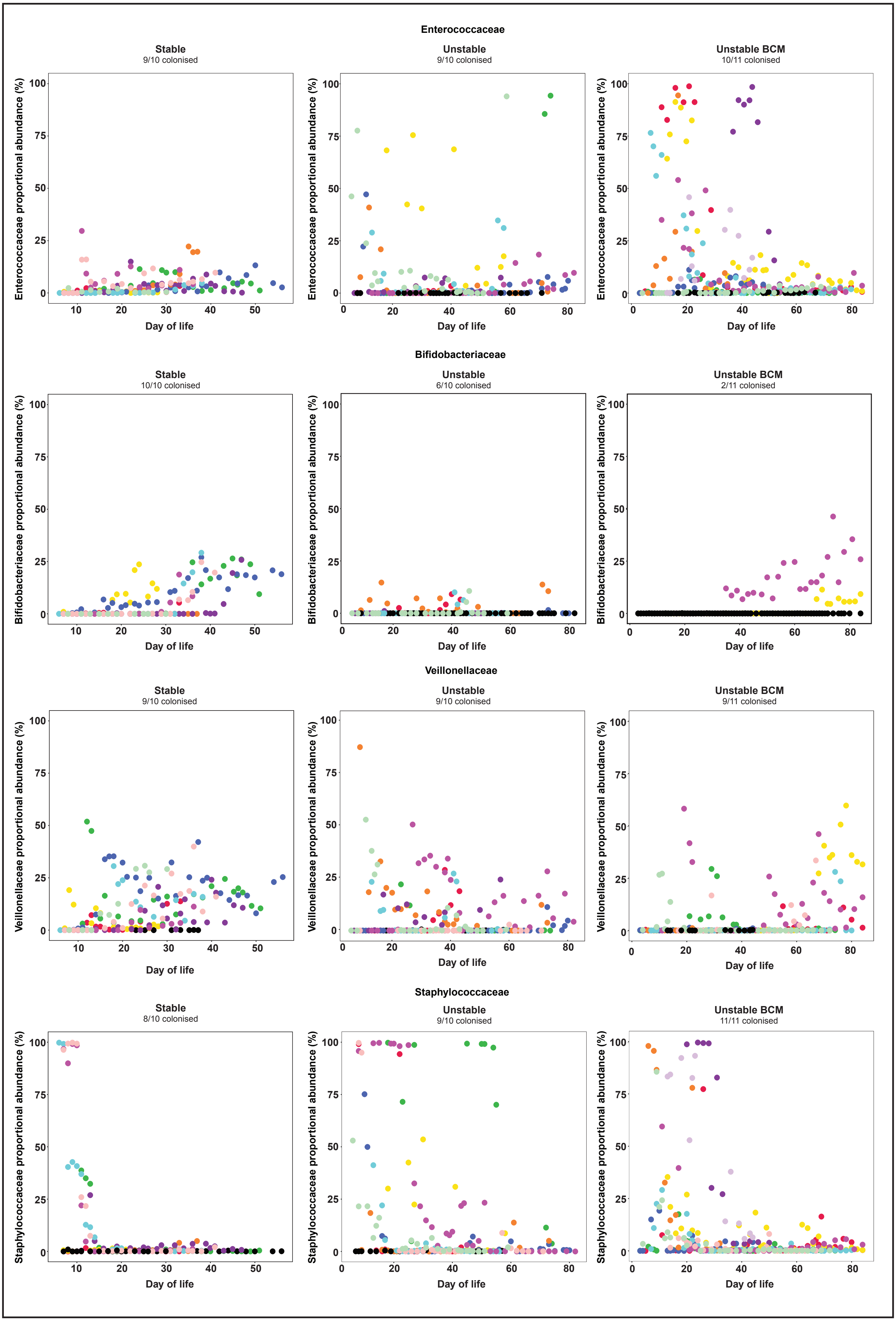

**Supplementary Figure 6**

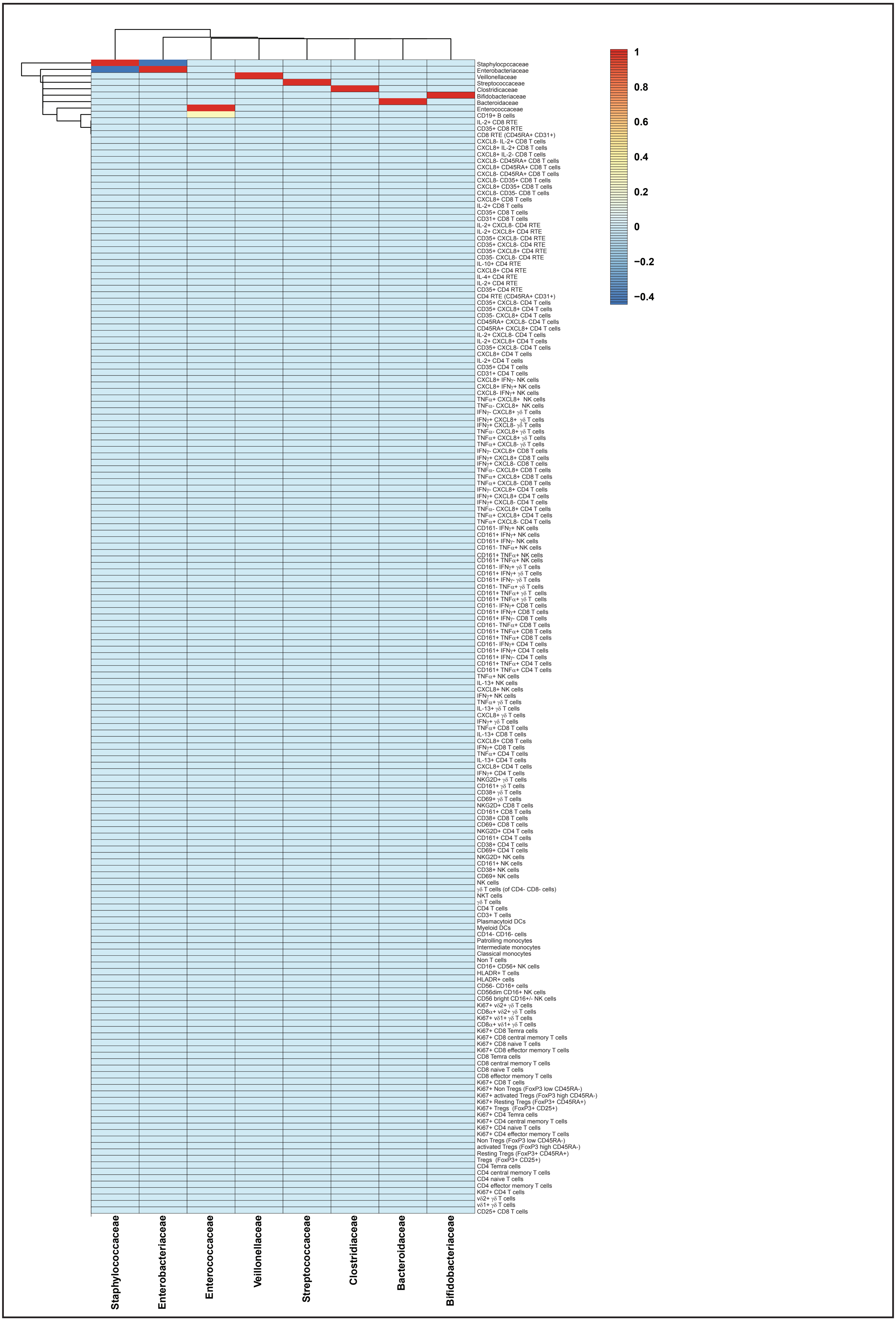

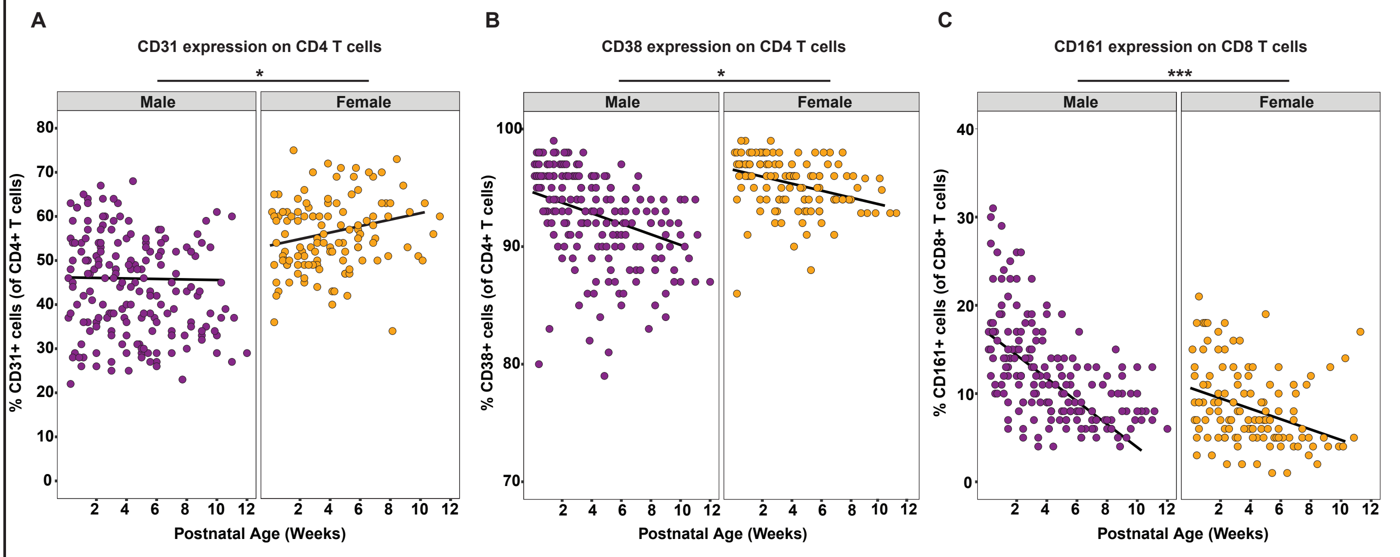

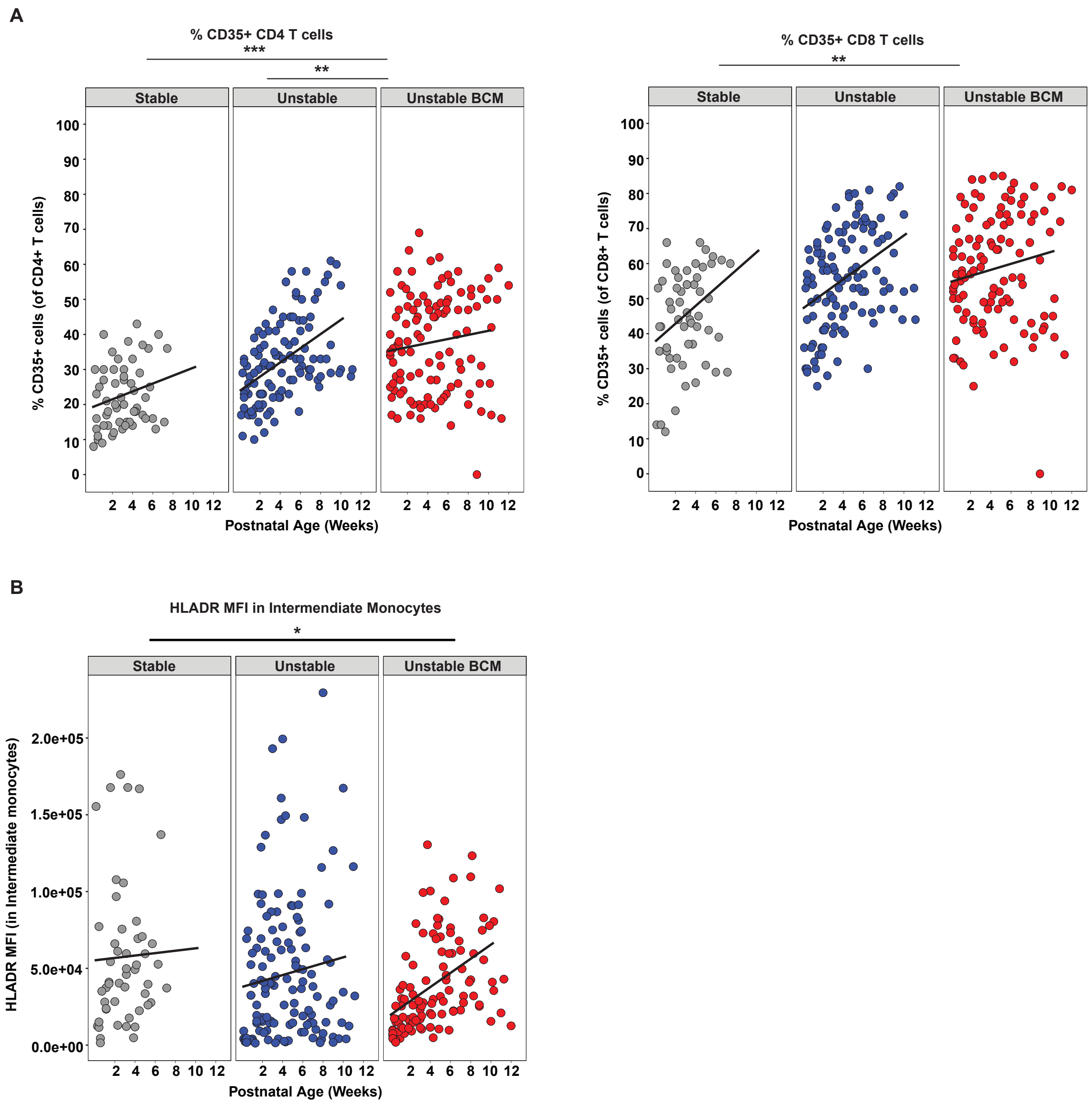

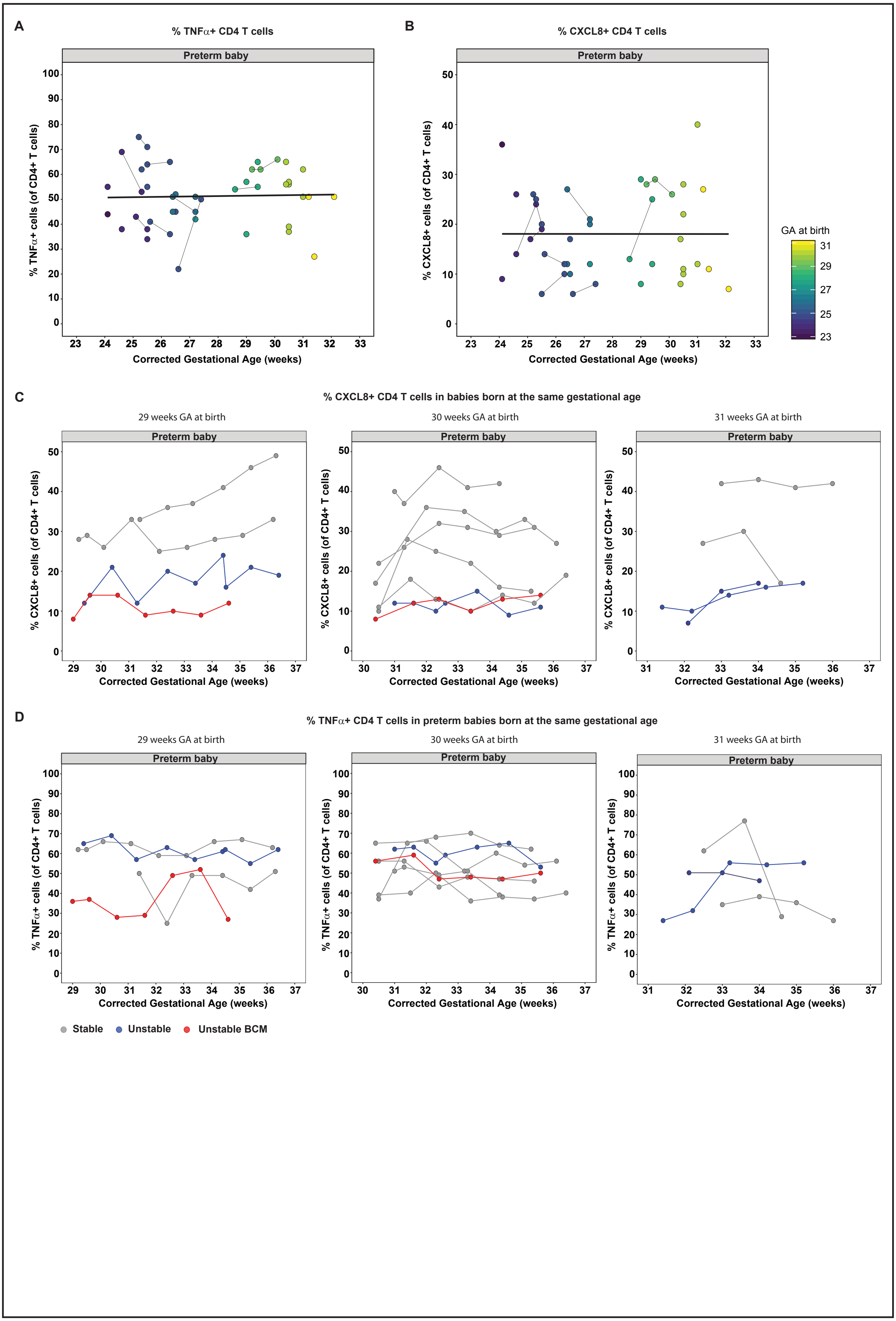
